# BMI1-mediated heterochromatinization represses G-quadruplex DNA formation to maintain genomic stability during replication

**DOI:** 10.1101/2024.12.23.630044

**Authors:** Roy Hanna, Gilbert Bernier

## Abstract

Single-stranded DNA secondary structures such as G-quadruplexes (G4s) can potentially disrupt transcription, replication, and repair. Using bio-informatic analysis, here we show that BMI1 is enriched at putative G4s flanked by heterochromatin domains, and that *BMI1* knockdown in human dermal fibroblasts (HDFs) resulted in heterochromatin relaxation and G4 induction, followed by replication stress and genomic instability. In these cells, G4s co-localized with large 53BP1 and PCNA foci resembling replication catastrophes. Inhibiting transcription partly attenuated DNA damage, suggesting rescue of transcription-replication collisions at difficult-to-replicate sequences. In *BMI1* knockdown or pyridostatin-exposed HDFs, the Werner helicase accumulated and co-localized with G4s. Acute *WRN* knockdown also resulted in G4 induction. In HDFs from Werner and Hutchinson-Gilford progeria syndromes, loss of heterochromatin and nuclear envelope anomalies were associated with G4 induction and DNA damage. Nuclear envelope anomalies were also prominent following *BMI1* knockdown. These findings suggests that heterochromatin-mediated repression of G4s attenuates replication stress and genomic instability, and that this mechanism is shared across distinct progeroid models.

## Introduction

Genome stability is essential for cellular proliferation during development and to sustain organ regeneration in adult life. For this, efficient mechanisms exist to ensure integrity of the replicated DNA strand and proper segregation of sister chromatids. When compromised, genomic stability can result in abnormal development, cancers and/or accelerated aging ^1^. The mammalian genome is subdivided in multiple sub-compartments, depending on the DNA sequence composition and localization on chromosomes ^2^. Euchromatin encompass transcriptionally active, gene-rich regions that are labeled by marks of open chromatin. Facultative heterochromatin is composed of tissue-specific and developmental genes labeled by the H3K27me3 and H2Aub histone marks. Constitutive heterochromatin contains repeat DNA sequences located at centromeric, pericentromeric and telomeric regions and that are labeled by the H3K9me3 histone mark. Intragenic and intergenic sequences, such as ALU repeat, Endogenous Retro-Viral element (ERV) and Long Interspersed Nuclear Element-1 (LINE-1 or L1), are also interspersed in the genome and transcriptionally silenced by deposition of the H3K9me3 histone mark and heterochromatinization ^2–4^.

Polycomb group proteins form large multimeric complexes involved in gene silencing through histone modification and chromatin compaction ^5,6^. The Polycomb repressive complex 2 is composed of EZH2, EED and SUZ12 and mediates H3K27me3 deposition ^5,7^. BMI1 (B-cell specific Moloney murine leukemia virus integration site 1), also called PCGF4, is a component of the canonical Polycomb repressive complex 1, which maintains chromatin compaction and developmental gene repression at facultative heterochromatin through its E3-mono-ubiquitin ligase activity mediated by RING1/RNF2 on histone H2A at lysine 119 (H2Aub) ^5,7^. *Bmi1^-/-^* mice present reduced post-natal growth and lifespan together with cerebellar degeneration ^8^. Bmi1 is important for somatic stem cells proliferation in part through inhibition of senescence-associated genes ^9^. Bmi1 also inhibits mitochondrial oxidative stress and *Bmi1^-/-^* mice show features of premature aging ^10,11^. In addition to the facultative heterochromatin, BMI1 is enriched at constitutive heterochromatin in somatic cells and co-purifies with architectural heterochromatin proteins ^12^. BMI1 inactivation in mice or in human cells causes loss of heterochromatin, heterochromatic genome instability, and reactivation of repetitive sequences ^12–14^. Cells deficient for BMI1 present chromosomal genomic instability ^15^. BMI1 plays multiple roles in DNA damage response (DDR) and repair, promoting recruitment of the DDR machinery and mono-ubiquitination of histones H2A and γH2AX at break sites, and homologous recombination and non-homologous end joining ^11,16–20^.

DNA sequences rich in tandemly spaced guanine quartets and capable of forming stable non-beta DNA secondary structures called G-quadruplexes (G4s) ^21–23^. G4s can potentially interfere with transcription, replication, and repair. Notably, it was shown that BMI1 or ATRX inactivation in human cells causes excessive formation of G4s, and that heterochromatin compaction was the main mechanism of G4 repression ^24,25^. By chromatin immunoprecipitation and sequencing (ChIP-seq) using the 1H6 antibody, it was found that 95% of G4s observed in human neurons originate from the transcription of evolutionary conserved L1s ^24^. L1s are 7k base pair retrotransposons representing nearly 17% of the entire human genome. L1s can be transcribed by RNAPol2 from a 5’ UTR promoter ^4,26,27^. However, less than 5000 evolutionary conserved L1 sequences have an intact internal promoter in the human genome, allowing for transcription ^4,26,27^. L1s contain multiple G4 motifs in their sequence, and the 3’ UTR of L1s was shown to form G4 structures *in vitro*, which could be important for propagation by retro-transposition ^24,28^.

In post-mitotic neurons, intragenic and intergenic accumulation of G4s was shown to interfere with gene transcription and mRNA splicing ^24^. In mitotic cells however, stable G4 structures can create an obstacle that stalls or slows down the replication fork, leading to replication stress, DNA recombination and genomic instability ^29–33^. Using human dermal fibroblasts (HDFs), we found here that acute *BMI1* knockdown results in loss of heterochromatin and G4 induction, which are rapidly followed by replication-associated DNA damage resembling mitotic catastrophes. Co-localization studies in *BMI1* knockdown cells further revealed a strong correlation between G4s and large DNA damage foci linked to replication units. Similarly, HDFs from Werner and Hutchinson-Gilford progeria (HGP) syndromes display a G4 phenotype associated with loss of heterochromatin. We conclude that in in replicating cells, heterochromatin is required to prevent excessive formation of G4s, which otherwise imped fork progression, potentially leading to mitotic catastrophes or genomic instability.

## Results

Using BMI1 ChIP-seq data from human neural progenitor (hNPC) cells, we identified 1366 BMI1 ChIP-seq peaks containing a putative G4 sequence ^34^. When we plotted the enrichment of H3K9me3 obtained from H3K9me3 ChIP-seq data around these peaks, the profile revealed the presence of 4 clusters, each showing unique H3K9me3 density at putative G4 sequences enriched for BMI1 (**Fig. 1A**). Clusters 1 and 4 showed the strongest H3K9me3 enrichment, with cluster 2 considered as negligeable because comprising less than 1% of the ensemble (**Fig. 1A-B**). Analysis of mean H3K9me3 density at putative G4 having a BMI1 peaks revealed a common distribution pattern for clusters 1 and 4 called peak-valley-peak pattern, with increasing H3K9me3 density on each side of the putative G4 sequence (**Fig. 1C**). This pattern is typical for histone marker deposition at regulatory elements ^35^. This pattern was not observed for cluster 2 and 3. It has been show that SIRT6 is enriched at the 5’ UTR of L1s, leading to mono-ADP ribosylation of KAP1 and heterochromatin-mediated repression of L1s transcription ^36^. We tested this possibility for BMI1 and found strong enrichment of BMI1 peaks at both the 5’ and 3’ UTRs of the evolutionary conserved L1H family of L1s (**Fig. 1D**). This revealed a strong correlation between BMI1 enrichment and H3K9me3 density at putative G4s and at L1s. Chromatin compaction was proposed as the main mechanism of G4 repression ^24,25^. To study G4 dynamics in replicating in HDFs, we first confirmed that BMI1 knockdown using a shRNA (shBMI1) resulted in lower H3K9me3 levels and induction of G4 at 16 hour (hr) post-transfection (**Fig. S1A**). To evaluate if inhibition of transcription and replication could affect G4s, HDFs transfected with shScramble (shScr) or shBMI1 plasmids were exposed or not to a high concentration of Aphidicolin, a DNA Polymerase alpha and delta inhibitor ^37^, and to DRB, an RNA Pol2 inhibitor^38^. BMI1 knockdown resulted in G4 induction, elevation of H3K9 acetylated (H3K9ac) levels – a mark of relaxed chromatin-, and downregulation of H3K9me3 (**Figs. 1E and S1B**). Notably, blocking transcription and replication nearly abolished G4 induction in shBMI1 HDFs while having little effect on chromatin state (**Figs. 1E and S1B**). This revealed that G4 induction can be uncoupled from chromatin state in replicating human cells.

**Figure 1.**
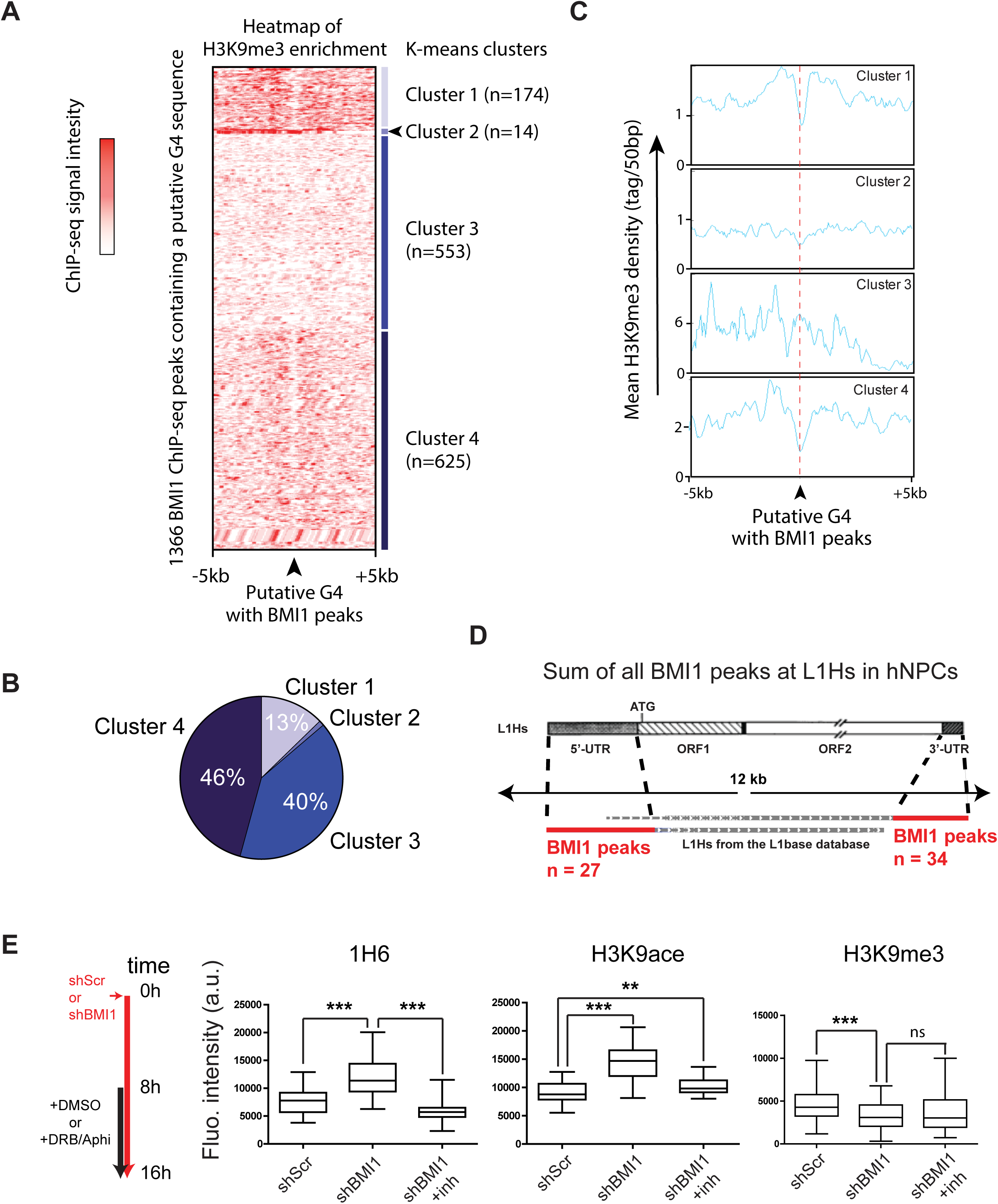
BMI1 is enriched at putative G4s and at high-density H3K9me3 domains. A. Representation of BMI1 ChIP-seq analysis, showing the distribution of BMI1 peaks containing putative G4 sequences, and along heatmap of H3K9me3 density. This revealed the presence of four clusters in human neural progenitor cells (hNPCs). B. Pie chart showing the relative proportion of the 4 clusters. C. Diagram showing the distribution of H3K9me3 domains along putative G4 sequences with BMI1 ChIP-seq peaks. Not the peak-valley-peak distribution of H3K9me3 around the G4. D. BMI1 ChIP-seq analysis in hNPCs showing enrichment of BMI1 peaks at the 5’ and 3’ of L1H sequences. E. Quantification of immunofluorescence experiments showing that G4 levels can be uncoupled from chromatin state, as revealed using transcription (DRB) and replication (Aphi) inhibitors in shBMI1 HDFs. N = 3 slides/condition and 200 cells/slide. Statistical differences were analyzed using an unpaired T-test with two tails. *P* ≤0.01**, *P* ≤0.001***. All values are means ± SEM.

To distinguish between transcription and replication, shBMI1 HDFs were exposed or not to DRB, Aphidicolin, or both at 8 hr, and EdU was added at 15:30 hr to label nascent DNA during S-phase (thus 30 minutes before analysis) (**Fig. 2A**). We first observed that EdU-positive cells were significantly less abundant in shBMI1 than in shScr HDFs, suggesting replication arrest (**Fig. 2B**). This difference remained even when transcription was inhibited (**Fig. 2B**). In second, we found that blocking transcription (or transcription and replication) in EdU-negative cells strongly reduced G4 levels (**Fig. 2A and C**, **white arrows in A).** This suggested that G4 induction in quiescent (G0/G1) shBMI1 cells is dependent on transcription, as previously reported in Alzheimer’s neurons ^24^. In contrast, blocking transcription in EdU-positive shBMI1 cells had no effect on G4 levels (**Fig. 2A and D**). This suggested that during S-phase, G4 induction in shBMI1 HDFs is largely dependent on replication. To further investigate this, we performed co-localization studies in cells exposed to EdU for 8 hr. We observed that 1H6 foci frequently co-localized with or were aligned close to EdU-positive foci in shBMI1 HDFs (**Fig. 2E**). The proliferating cell nuclear antigen (PCNA) is a scaffold for proteins at the replication fork ^39,40^. We found that 1H6 and PCNA generally did not co-localized but showed close association in shBMI1 HDFs, with two 1H6 foci frequently bordering a single PCNA foci (**Fig. 2E**). This revealed a close association between G4s and the DNA replication machinery in BMI1 knockdown cells.

**Figure 2.**
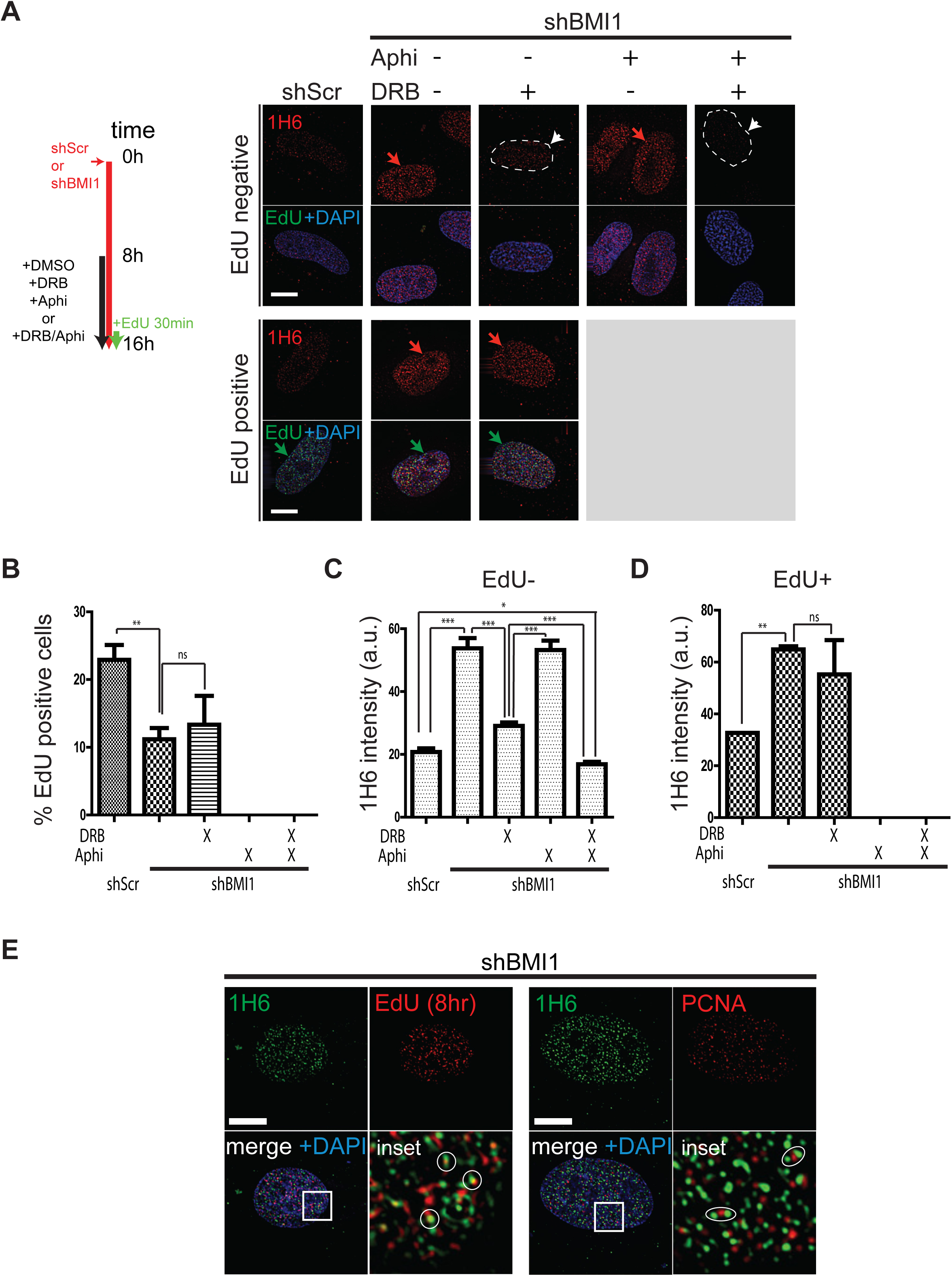
G4 induction in shBMI1 HDFs depends on transcription in quiescent cells. A. Immunofluorescence experiments of shScr vs drug-treated or untreated shBMI1 HDFs. Drugs were added at 8hr, and EdU was added at 15h30hr (30min). Quiescent (EdU negative) shBMI1 cells show dependency on transcription, or transcription and replication, for G4 induction (dashed line-white arrows). Replicating (EdU positive-green arrows) shBMI1 cells show independency on transcription for G4 induction. Red arrows point to 1H6 (G4) foci. Scale bar: 5 µm. B. Quantification of experiments in (A) showing reduced number of EdU+ cells in shBMI1. C. Quantification of experiments in (A) showing reduced 1H6 intensity in EdU-shBMI1 HDFs exposed to DRB. D. Quantification of experiments in (A) showing that DRB does not affect 1H6 intensity in EdU+ shBMI1 HDFs. (C-D) N = 3 slides/condition and 200 cells/slide. Statistical differences were analyzed using an unpaired T-test with two tails. *P* ≤0.01**, *P* ≤0.001***. All values are means ± SEM. E. Immunofluorescence analysis of shBMI1 HDFs at 16Hr showing close association between G4s and EdU (8 hours labeling) or PCNA, as shown in the insets. Scale bar: 5 µm.

To test the possibility that G4s could be induce by replication stress, we exposed shScr or shBMI1 HDFs to low concentration of the DNA replication inhibitors Aphidicolin or to Hydroxyurea. Hydroxyurea is a ribonucleotide diphosphate reductase inhibitor causing depletion of deoxyribonucleotides ^41^. When exposed to Aphidicolin or Hydroxyurea and analyzed 16hr post-transfection, both shScr and shBMI1 HDFs showed an increased in G4s when compared to media-only, with Aphidicolin showing the strongest effect (**Fig. 3A**). These findings revealed that mild replication stress, which promotes new replication origins and formation of single strand DNA (ssDNA), can induce G4s in normal cells and exacerbate the G4 phenotype in BMI1-deficiency cells ^42^. They also suggest an intimate interconnection between G4s and replication origins, which are highly enriched in G4 motifs ^43^. Under replicative stress conditions, non-beta DNA structures such as G4s can interfere with fork elongation, leading to DNA breaks ^44,45^. Replication stress in general can also result into DNA damage ^42,46^. To test if G4s were associated with DNA damage, we performed a time course analysis to evaluate G4s and DNA damage in shBMI1 HDFs. 53BP1 is a generic marker of DNA damage foci ^47^. 53BP1 also labels nuclear bodies under replication stress conditions ^46,48^. At time 0 hr, G4s and 53BP1 foci were near absent in shBMI1 HDFs (**Fig. 3B**). At 4 hr, G4s were readily visible, and large DNA damage foci positive for 53BP1 were apparent at 16 hr (**Fig. 3B-white arrow**). To investigate this, we compared the number and size of 53BP1 foci between shScr and shBMI1 HDFs. BMI1-knockdown HDFs presented significantly more average size (0.5-1.5 μm^2^ in diameter) and large size (<1.5 μm^2^ in diameter) 53BP1-positive foci compared to shScr (**Fig. 3C**). EdU labeling of shBMI1 HDFs for 8 hr or 30 minutes before fixation showed a significant reduction of EdU-positive cells having large 53BP1 foci at 30 minutes, suggesting that large 53BP1 foci generated during S-phase induce cell cycle arrest (**Fig. S2A-B**). We next exposed shBMI1 HDFs to DRB, Aphidicolin, or DRB + Aphidicolin 8 hr post-transfection. We found that when compared to DMSO, all conditions prevented apparition of large 53BP1 foci, but that average size DNA damage foci partly remained when cells were only exposed to DRB (**Fig. 3D**). This suggested that in shBMI1 HDFs, both large and medium size DNA damage foci are transcription and replication dependent, and that collisions between the transcription and replication machinery may be involved ^18^.

**Figure 3.**
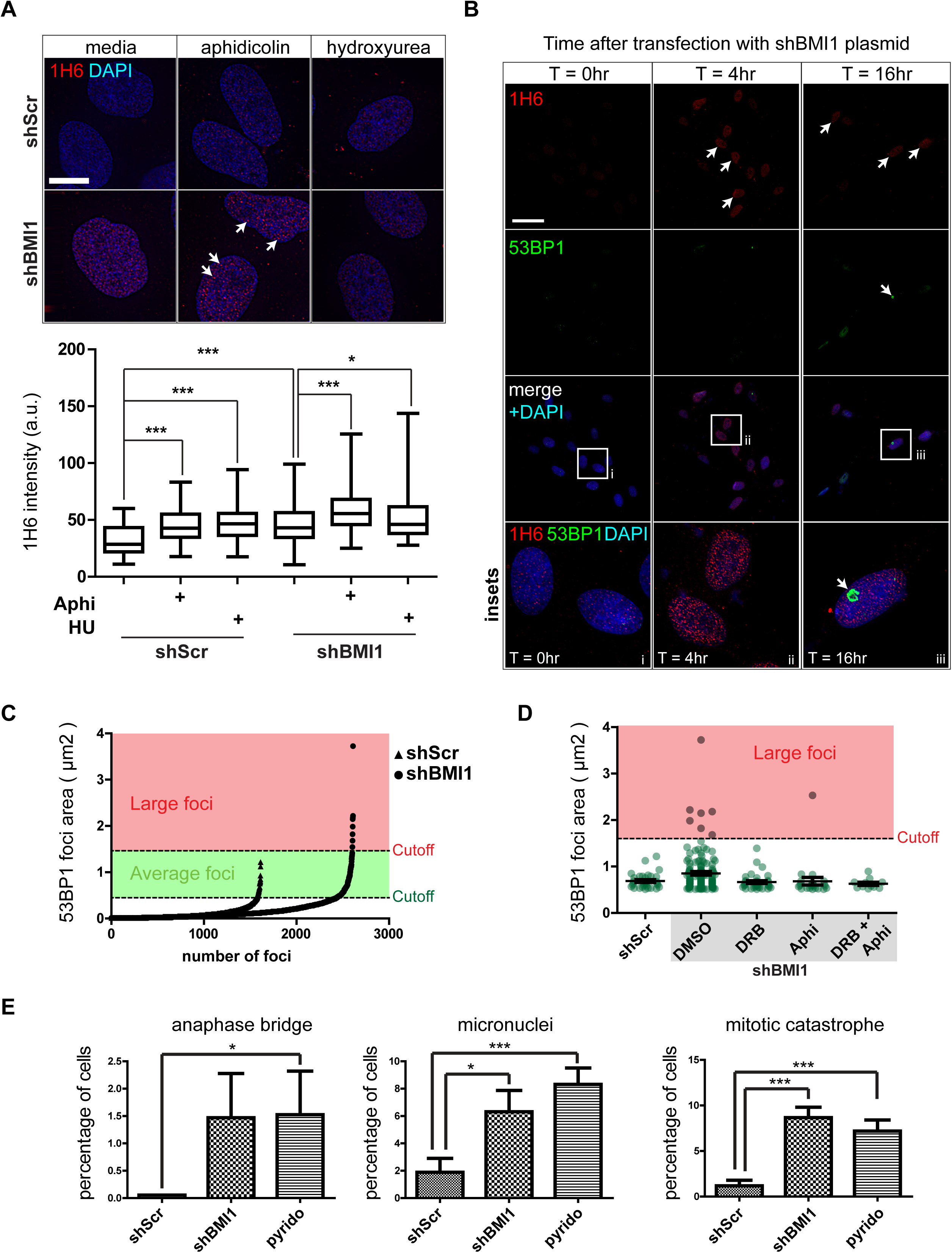
BMI1 knockdown in HDFs induces genomic instability. A. Immunofluorescence analysis of HDFs at 24hr post-transfection with shScramble or shBMI1 plasmids. HDFs were exposed or not exposed to low concentration of the DNA replication stress agents aphidicolin or hydroxyurea for 8hr before fixation. G4 structures were labeled with the 1H6 antibody (white arrows). Scale bar: 10 µm. Bottom panel: Quantification of the experiments. 3 slides/condition and 200 cells/slide. Statistical differences were analyzed using an unpaired T-test with two tails. *P* ≤0.05*, *P* ≤0.001***. All values are means ± SEM. B. Time course analysis of HDFs knockdown for BMI1 and visualized by immunofluorescence. Note that G4 structures (white arrows) are induced before 53BP1. A large DNA damage foci labeled with 53BP1 is showed in the inset (white arrow). Scale bar: 10 µm. C. Histogram showing the size distribution of 53BP1 DNA damage foci 24hr after transfection of shScramble or shBMI1 plasmids in HDFs. DDR foci were considered as large when bigger that 1.5 scare mm. Cutoff was used to define the minimal size of a DDR foci. D. Histogram showing the size distribution of 53BP1 DNA damage foci 24hr after transfection of shScramble or shBMI1 plasmids in HDFs and treated or not treated for 8hr with DMSO, DRB and/or aphidicolin. E. Histograms showing the frequency of anaphase bridge, micronuclei and mitotic catastrophe 72 hours after transfection of shScramble or shBMI1 plasmids in HDFs. Those were compared to HDFs exposed to pyridostatin for 72 hours. 3 slides/condition and 200 cells/slide. Statistical differences were analyzed using an unpaired T-test with two tails. *P* ≤0.05*, *P* ≤0.001***. All values are means ± SEM.

To further probe for a possible link between transcription and DNA damage, we compared shScr and shBMI1 HDFs with HDFs treated with pyridostatin or exposed to 10Gy ^49^. Co-localization studies showed an absence of correlation between RNA Pol2 and 53BP1 foci in all conditions except for BMI1 knockdown HDFs, which showed a modest but significant R coefficient of 0.15 (**Fig. S2C**). Because RNA Pol2 foci were much more abundant than 53BP1 foci, we quantified the proportion of 53BP1 that were also positive for RNAPol2. This revealed that ∼75% of 53BP1 foci in BMI1 knockdown HDFs were positive for RNA Pol2, compared to ∼20% for CTL HDFs irradiated with 10Gy (**Fig. S2D**). Anaphase bridges, micronuclei and mitotic catastrophes are common consequences of replication stress ^42^. Pyridostatin is a G4 stabilizer compound that can induce replication stress ^49,50^. When compared to shScr, we observed that after 72 hrs, both shBMI1 or control HDFs exposed to pyridostatin showed genomic instability as revealed by the large number of anaphase bridges, micronuclei and mitotic catastrophes (**Fig. 3E**). These results are consistent with previous work showing chromosome breaks in immortalized and normal hematopoietic cells deficient for BMI1 ^15^. They also show that DNA damage is not the driving factor behind G4 structure accumulation. Instead, G4 structures formed as a result of chromatin relaxation and next appeared to drive DNA damage. To investigate this possibility, we performed co-localization studies. When compared to controls, shBMI1 HDFs showed a highly significant R coefficient of 0.58 between G4s and 53BP1, with robust co-localization between G4s and large DNA damage foci (**Fig. 4A-white arrows**). Co-localization studies also confirmed the strong correlation between 53BP1 and γH2AX foci, and between 53BP1 and PCNA foci (**Fig. 4B and C-white arrows**) ^51^. This revealed strong association between G4s and large DNA damage foci at replication units in BMI1 knockdown HDFs.

**Figure 4.**
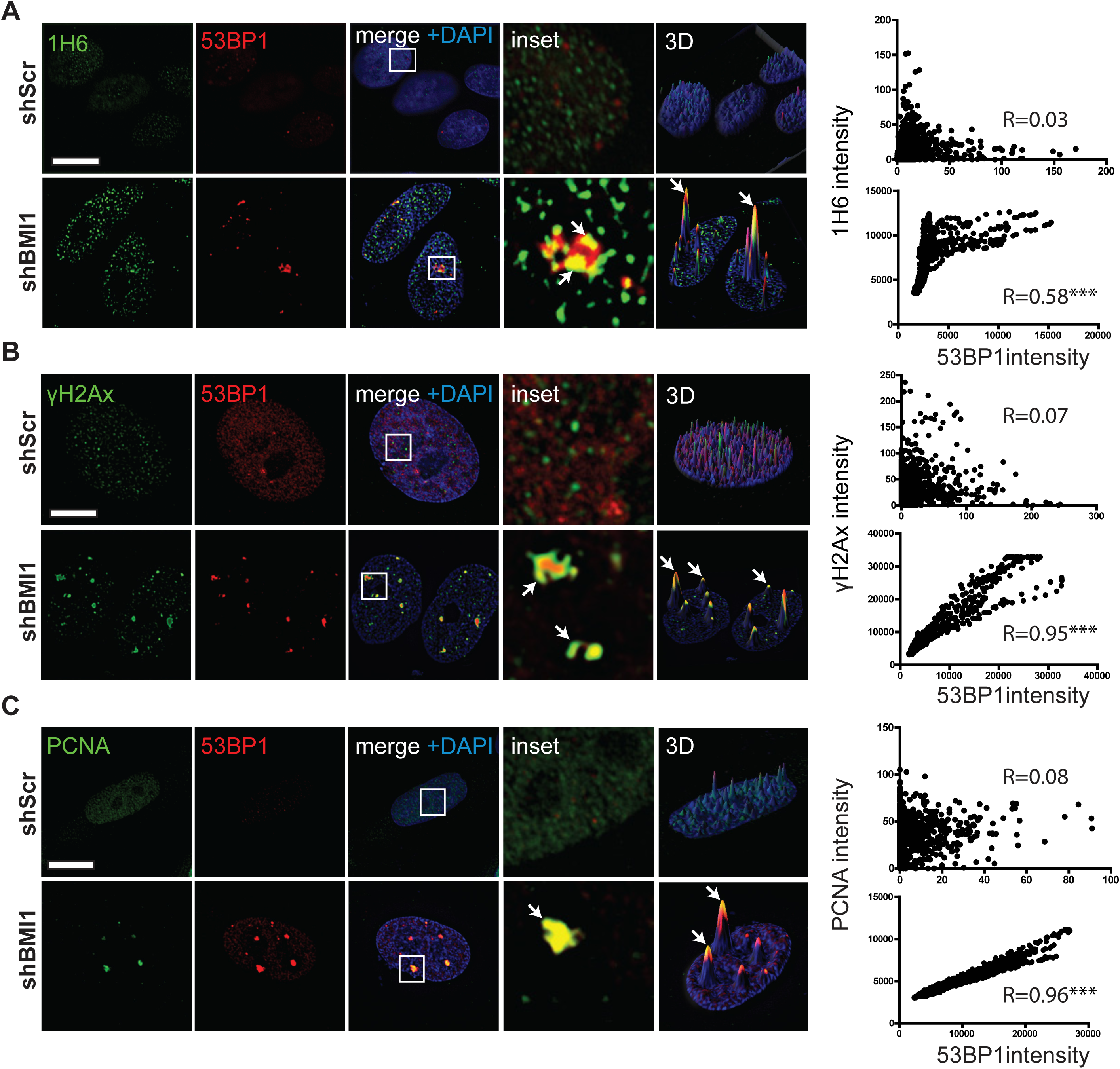
G4s associate with DNA damage at replication units following BMI1 knockdown. A-C. Immunofluorescence analysis of HDFs at 24hr post-transfection with shScramble or shBMI1 plasmids. Scale bar: 5 µm. A. Labeling with 1H6 and 53BP1 antibodies showing co-localization (arrows) in shBMI1, with R = 0.58. B. Labeling with γH2AX and 53BP1 antibodies showing co-localization (arrows) in shBMI1, with R = 0.95. C. Labeling with PCNA and 53BP1 antibodies showing co-localization (arrows) in shBMI1, with R = 0.96. N = 200 cells/group. Statistical differences were analyzed using an unpaired T-test with two tails. *P* ≤0.001***. All values are means ± SEM.

The ataxia telangiectasia and Rad3-related kinase ATR is essential for the maintenance of genomic stability and is activated by single-stranded DNA (ssDNA) ^52^. The presence of ssDNA is common at stalled replication forks and as intermediate during DNA repair ^52^. Genome-wide studies also identified structured DNA and short tandem-repeats as principal sites of fork collapse when ATR is inhibited ^53^. Using immuno-fluorescence, we observed low p-ATR levels in shScr HDFs. In contrast, p-ATR levels were high in both shBMI1 and pyridostatin-treated HDFs at 24 hr, showing partial co-localization with G4s using the 1H6 antibody (**Fig. 5A**). These results suggested replicative stress in shBMI1 and pyridostatin-treated HDFs. Werner syndrome is rare progeroid genetic disease linked to mutations in *WRN* ^54^. *WRN* encodes an RecQ like helicase that can recognize and resolve non-beta DNA secondary structures ^55,56^. WRN is also implicated in the recovery of stalled replication forks and is directly phosphorylated by ATR, allowing proper WRN localization at RPA-foci ^57^. In shScr HDFs, WRN levels were relatively low. In both shBMI1 and pyridostatin-treated HDFs, WRN was highly induced in the nucleus and showed near perfect co-localization with 1H6 (**Fig. 5B**).

**Figure 5.**
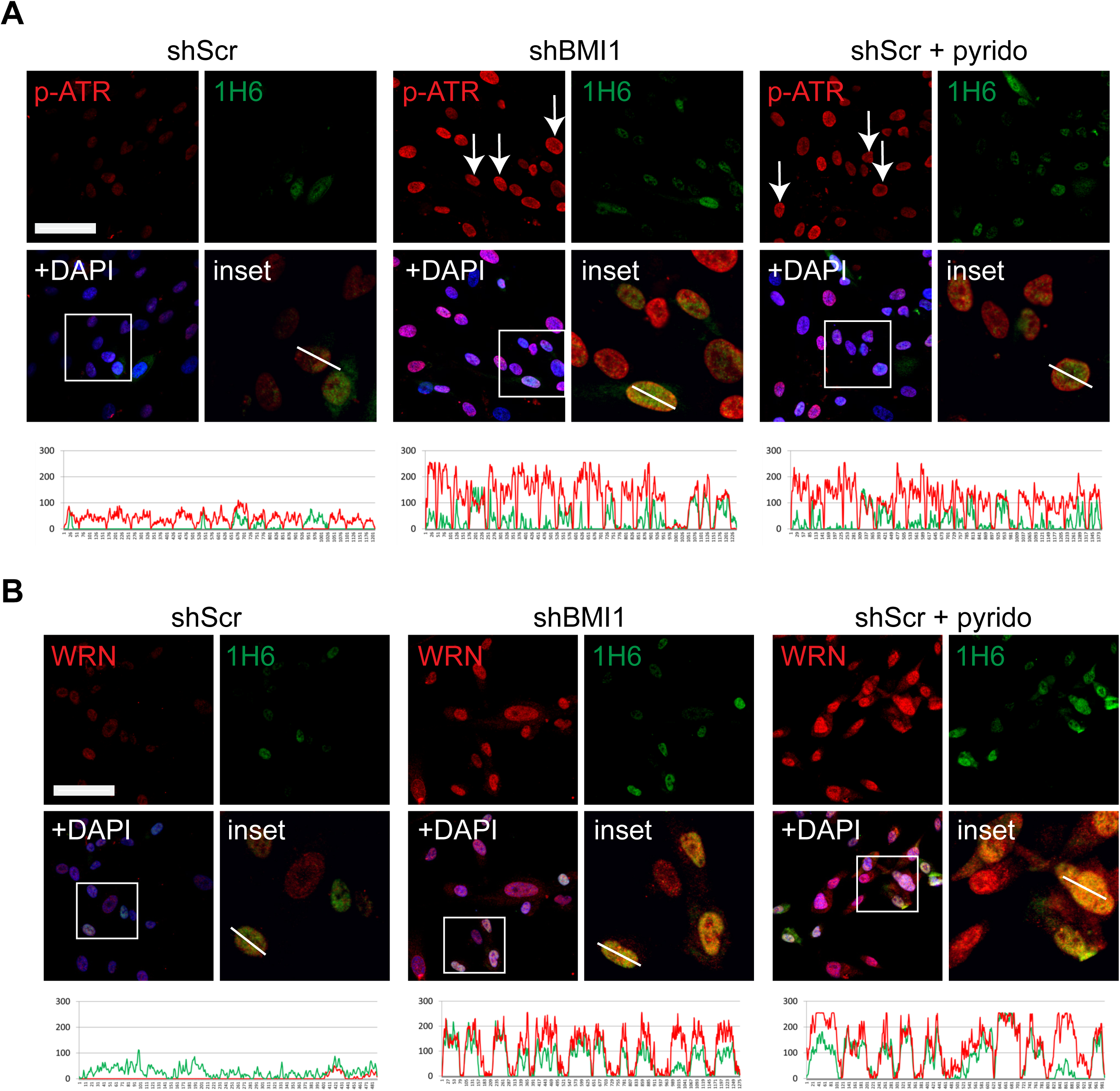
p-ATR and WRN accumulate in BMI1 knockdown HDFs. A. Immunofluorescence analysis of HDFs at 24hr post-transfection with shScramble or shBMI1 plasmids and in shScramble HDFs treated with pyridostatin (pyrido). Scale bar: 50 µm. Note the robust induction of p-ATR in shBMI1 and pyridostatin-treated fibroblasts (arrows). Histograms on the bottom show quantification analysis of p-ATR (in red) and 1H6 (in green). B. Immunofluorescence analysis of HDFs at 24hr post-transfection with shScramble or shBMI1 plasmids and in shScramble HDFs treated with pyridostatin (pyrido). Scale bar: 50 µm. Note the robust induction of WRN in shBMI1 and pyridostatin-treated fibroblasts (arrows). Histograms on the bottom show quantification analysis of WRN (in red) and 1H6 (in green). Note the robust co-localization between WRN and G4s (1H6) in shBMI1 and pyridostatin-treated cells.

To evaluate the speed of WRN recruitment following G4 stabilization, we performed a time course study. Upon pyridostatin exposure, we found that G4 accumulation was readily visible in the nucleus after 15 minutes using the 1H6 antibody, while WRN accumulation rapidly followed after 30 minutes (**Fig. 6A**). Conversely, acute *WRN* knockdown in HDFs using siRNA resulted in robust G4 induction and apparent nuclear envelope deformation after only 24 hrs (**Fig. 6B**). This suggested that WRN is rapidly recruited to newly forming G4s and that *WRN* is required to prevent G4 accumulation, which is consistent with WRN RecQ like helicase activity ^55,56^. Loss of heterochromatin and nuclear envelope anomalies are important features of cells from both Werner and HGP syndromes ^54,58^. HGP is linked to a dominant point mutation in *LMNA* encoding for LaminA/C, and thus unrelated to G4 unwinding activity. In contrast, *WRN* mutation may affect both heterochromatin-mediated repression of G4s and WRN-mediated resolution of G4s. To compare these, we investigated HDFs from Werner and HGP syndromes. As expected, Werner and HGP fibroblasts presented heterochromatin and nuclear envelope anomalies (**Fig. 6C-D**). Notably, they also both showed robust G4 induction and accumulation of DNA damage (**Fig. 6C-D**). This suggested that loss of heterochromatin may be the common mechanism leading to G4 induction in Werner, HGP and shBMI1 fibroblasts. It was proposed that resemblance between Werner and HGP phenotypes could be explained by de-repression of *progerin* in human Werner cells ^59^. We thus investigate if *BMI1* knockdown also resulted into a laminopathy and in *progerin* expression. We labeled shBMI1 HDFs with LaminA/C at 24 hr post-transfection. We found that 74% of shBMI1 HDFs presented nuclear envelope anomalies, such as deformation and folding, compared to 29% in shScr HDFs, and that this difference was highly significant (**Fig. 7A**). Western blot and quantitative RT-PCR analyses revealed that *BMI1* knockdown was not associated with alterations in LaminA/C levels or ratio, nor with elevated expression of *progerin* (**Fig. 7B-C**) ^60^. It was however associated with elevation of *p16-INK4A* (also called *CDKN2A*) expression-a direct target of the BMI1/RING1B complex repressive activity ^61^. These results suggested the existence of a common pathological mechanism in progeroid cells, where loss of heterochromatin leads to G4 induction, nuclear envelope deformation, and replication-associated genomic instability.

**Figure 6.**
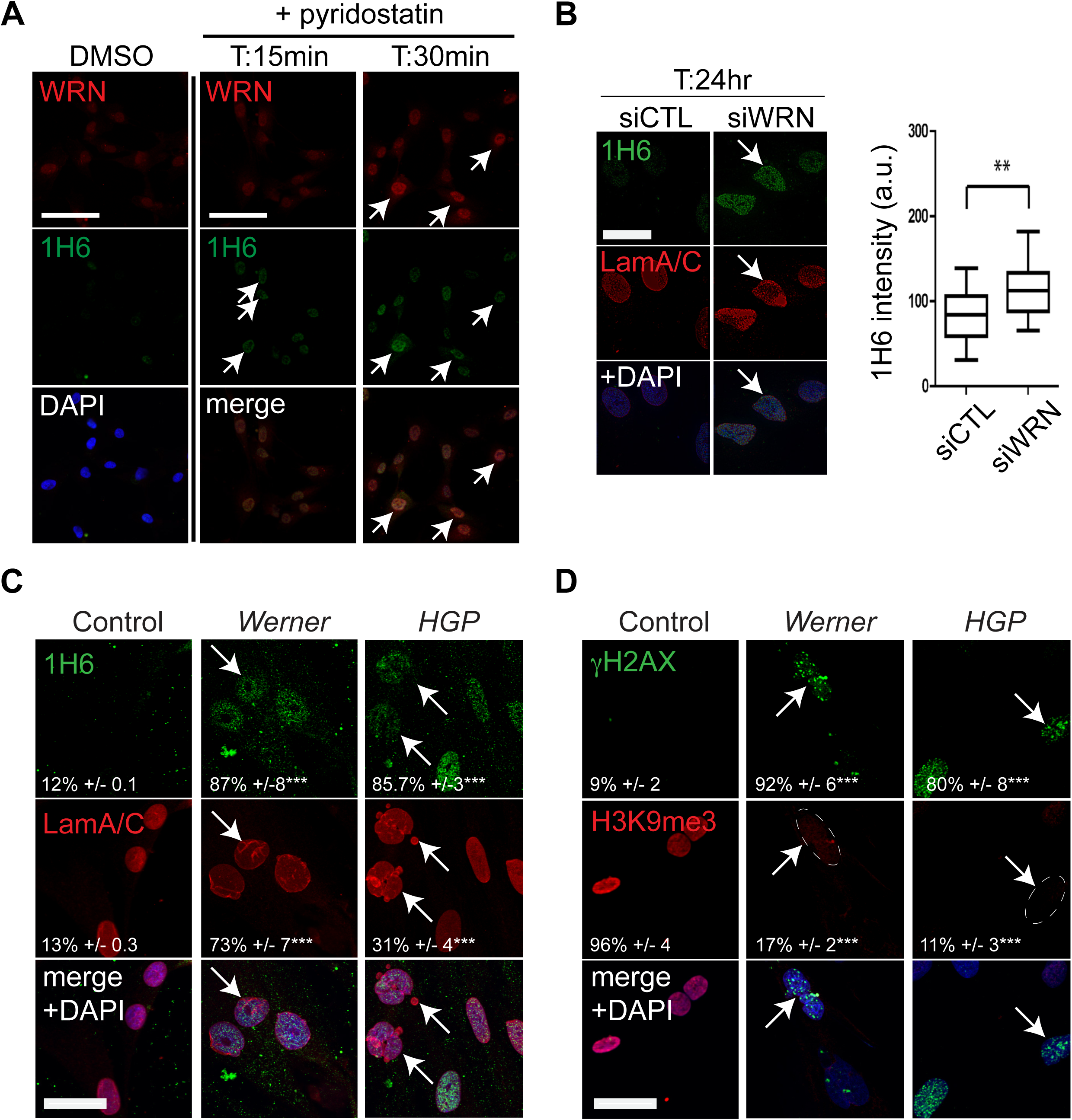
Loss of heterochromatin and G4 induction in Werner and HGP fibroblasts. A. Immunofluorescence analysis of HDFs treated with DMSO or pyridostatin. Note the rapid induction of G4s at 15min (arrows) and the accumulation of WRN at 30min (arrows). Scale bar: 50 µm. B. Immunofluorescence and quantitative analyses of HDFs at 24hr post-transfection with siCTL or siWRN oligonucleotides. Note the induction of G4s (1H6) and alteration of the nuclear envelope (Lamin A/C) in siWRN fibroblasts (arrows). (a.u.) arbitrary units. Scale bar: 10 µm. C-D. Immunofluorescence analysis of Werner and HGP fibroblasts, showing G4 (1H6) induction and nuclear envelope (Lamin A/C) alterations (arrows), together with DNA damage (γH2AX) and heterochromatin (H3K9me3) loss (arrows). Scale bar: 15 µm. N = 200 cells/group. Statistical differences were analyzed using an unpaired T-test with two tails. *P* ≤0.001***. All values are means ± SEM.

**Figure 7.**
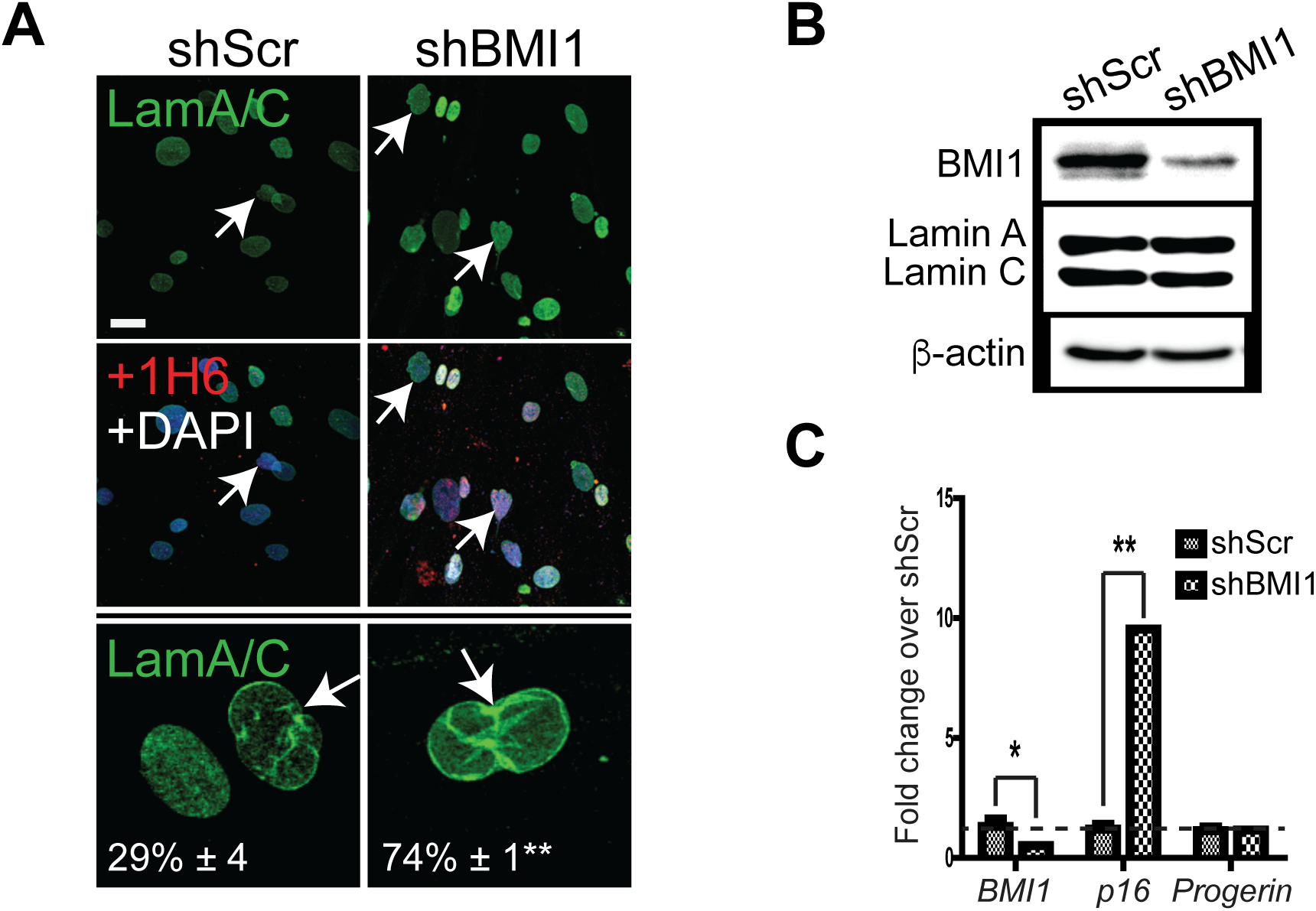
BMI1 knockdown in HDFs induces a laminopathy. A. Immunofluorescence analysis of HDFs at 24hr post-transfection with shScramble or shBMI1 plasmids. Scale bar: 10 µm. Deformation of the nuclear envelope, as labeled with Lamin A/C, was more frequent in shBMI1 HDFs (arrows). The bottom panel is a high magnification of cells with nuclear envelope deformation. N = 200 cells/group. Statistical differences were analyzed using an unpaired T-test with two tails. *P* ≤0.01**. All values are means ± SEM. B. Western blot analysis of HDFs at 24hr post-transfection with shScramble or shBMI1 plasmids. Note BMI1 reduction in shBMI1 cells. C. Real-time PCR analysis of HDFs at 24hr post-transfection with shScramble or shBMI1 plasmids. While *BMI1* expression was reduced and *P16INKA* was increased in shBMI1 cells, *progerin* levels were unaffected. Statistical differences were analyzed using an unpaired T-test with two tails. *P* ≤0.05*, and *P* ≤0.01**. All values are means ± SEM.

## Discussion

We showed here that in normal human replicating cells, BMI1 is enriched at putative G4s flanked by H3K9me3-labeled heterochromatin domains. *BMI1* knockdown in HDFs resulted in loss of H3K9me3 and rapid induction of G4s. In quiescent cells, G4 induction was dependent on transcription. In S-phase cells, G4 induction was dependent on replication. Large DNA damage foci observed in *BMI1* knockdown cells co-localized with G4s and the DNA replication machinery. Formation of these damage was attenuated by blocking transcription, correlating with partial co-localization of 53BP1 with RNAPol2. *BMI1* knockdown or pyridostatin treatment resulted in the accumulation of p-ATR and WRN. WRN co-localized with G4s and *WRN* knockdown resulted in G4 induction. HDFs from Werner and HGP syndromes showed loss of heterochromatin, G4 induction, DNA damage and laminopathy, suggesting a unifying mechanism linking progeroid syndromes.

We uncovered that G4s could be uncoupled from chromatin state in shBMI1 HDFs simultaneously exposed to DRB and Aphidicolin. This suggests that preventing transcription and replication, and thus ssDNA formation, is sufficient to prevent G4s. We have also found that BMI1 peaks are enriched at 5’ and 3’ UTRs of evolutionary conserved L1s, suggesting that BMI1 directly or indirectly represses L1 transcription^12,62–66^. Assuming that most G4s detected in HDFs using the 1H6 antibody originate from L1s, as in neurons, our work supports a model where heterochromatin-mediated inhibition of L1s transcription or replication is the primary mechanism preventing the formation of G4 structures in dividing cells. In this model, G4s can be generated in two ways: through excessive L1s transcription upstream of the replisome, and through L1s undergoing replication initiation. In both scenarios, G4s are predicted to represent an obstacle to fork progression. We found that blocking transcription cannot prevent G4s in *BMI1* knockdown cells during S-phase (**Fig. 2**). How this can be explained by loss of heterochromatin is unclear, since the replisome is by default free of nucleosomes ^67^. One possibility is that *BMI1* knockdown prevents normal recruitment of proteins having G4 helicase activity at the replisome. BMI1 co-purifies with ATRX, and ATRX was proposed to interact with MCM DNA helicases to resolve G4 structures on the newly replicated strand ^12,25^. It was also showed that Ezh2 inactivation in embryonic stem cells impairs H3K27me3 deposition near gene’s promoters, leading to an increase in the number of replication origins and of DNA replication initiation activity ^68^.

Interestingly, this anomaly was independent of transcription and presumably mediated by reduced nucleosome density ^68^. Herein, we found that G4s can be induced by a mild replication stress, which is known to activate new replication origins. In mammalian cells, replication origins are enriched in dimeric G4 motifs and activated in nucleosome-free regions ^43,69^. While speculative, loss of heterochromatin may promote the formation of new nucleosome-free regions, thus increasing the number of G4-rich replication origins and origin activity. We also established that blocking transcription in *BMI1* knockdown cells could attenuate replication-associated DNA damage. This is in agreement with previous findings showing that the BMI1/RNF2 complex, through mono-ubiquitination of histones H2A/H2AX and repression of RNAPol2 activity, helps preventing transcription-replication conflicts at difficult-to-replicate and repair regions ^16–18,70^. Our results thus suggest that following *BMI1* knockdown, loss of heterochromatin precedes induction of G4s, triggering replication stress and genomic instability.

Werner and HGP syndromes show different disease onset, severity and phenotypes ^54^. Both syndromes neither perturb brain development nor are associated with neurodegeneration. In contrast, *Bmi1*-null mice show reduced growth and lifespan, premature aging features, defective hematopoiesis, and degeneration of the brain and retina ^8–11,14^. Yet, despite the phenotypic difference between these 3 progeroid syndromes, several characteristics are shared between them at the cellular level. The heterochromatin loss model of aging stipulates that heterochromatin erosion promotes epigenomic and genomic instability, leading to cellular and organismal aging ^71,72^. In pioneer work, it was demonstrated that heterochromatin undergo erosion during aging and that perturbation of components of the NURD complex results into loss of heterochromatin which precedes DNA damage ^73^. We previously showed that ∼50% of the BMI1 protein pool was associated with the SDS-soluble constitutive heterochromatin fraction, and that BMI1 co-precipitated with architectural heterochromatin proteins ^12^. Interestingly, WRN protein can co-precipitate with SUV39H1, HP1α and LAP2β, suggesting a direct role in heterochromatin organization ^74^. While the mechanism leading to heterochromatin erosion in HGP cells is unknown, it is conceivable that loss of interaction between the nuclear envelope and lamina-associated chromatin regions is sufficient to trigger heterochromatin disorganization ^75,76^. Hence, we have found that loss of heterochromatin following *BMI1* knockdown rapidly impairs nuclear envelope integrity, suggesting that the maintenance of both structures is interdependent. Interestingly, progerin was shown to trigger DNA damage during DNA replication ^77^. DOX-inducible expression experiments further revealed that progerin induces loss of heterochromatin in G1-arrested cells without causing DNA damage, and that DNA damage only appear during late S-phase in cells lacking heterochromatin ^78^. These observations suggest that both *WRN* deficiency and progerin expression can directly affect heterochromatin stability. However, a unifying mechanism explaining S-phase associated genomic instability in Werner and HGP cells is lacking. We have shown that WRN is rapidly recruited to G4s, that *WRN* knockdown results in G4 induction, and that G4s accumulate in Werner and HGP fibroblasts. That leaves open the possibility that loss of heterochromatin in Werner and HGP cells results in the accumulation of G4s at replication forks, leading to replication stress and genomic instability.

It was shown that loss of heterochromatin can also result in de-repression of ERVs and L1s, leading to retro-transposition and activation of the cGAS/STING pathway ^4,36,79^. In various aging models, blocking retro-transposition with a reverse transcriptase inhibitor attenuates senescence and genomic instability ^1,79^. Thus, genomic instability resulting from loss of heterochromatin may originate from multiple sources i.e. accumulation of G4s and retro-transposition of L1s and ERVs. Whether blocking retro-transposition in *BMI1*-deficient cells or in *Bmi1*-null mice can attenuate DNA damage, cellular senescence, neurodegeneration and/or aging phenotypes should be tested.

### Limitations of the study

In this work, we have not demonstrated whether G4 induction correlated with fork elongation blockage or DNA damage formation during replication in Werner and HGP cells. Further work is thus required to establish a direct link between G4 induction and genomic instability in Werner and HGP cells.

## Acknowledgments

This work was supported by a grant from the National Science and Engineering Research Council of Canada (NSERC). R.H. was supported by fellowships from the Molecular Biology Program of Université de Montréal.

## Data availability statements

Raw data, cell lines, and reagents are available upon request.

## Code availability statement

None

## Author contributions

Conceived and designed: G.B. and R.H.

Performed the experiments: R.H.

Analyzed the data: G.B., R.H.

Wrote the paper: G.B., R.H.

## Materials and Methods

### Cell cultures

Normal human dermal fibroblasts (HDFs) were purchased from the Coriell Institute. HDFs were cultured with DMEM/F12 media (Invitrogen) supplemented with 10% FBS (Invitrogen) and non-essential amino acids (Invitrogen). Pyridostatin was used at a concentration of 5µM (Sigma, SML0678-5MG). For the replication and transcription arrest we used respectively: 1μg/mL of Aphidicolin from Nigrospora sphaerica (Sigma, A0781-1MG) and 40μM of 5,6-Dichlorobenzimidazole 1-β-D-ribofuranoside (DRB; Sigma, D1916-10MG). A concentration of 0.2μg/mL of aphidicolin or of 0.2mM of Hydroxyurea (Sigma, H8627) was used to induce replication stress.

### Immunofluorescence

Eyes were extracted and fixed by immersion over-night at 4°C in 4% paraformaldehyde (PFA)/3% sucrose in PBS, pH 7.4. Samples were washed three times in PBS, cryoprotected in PBS/30% sucrose, and frozen in CRYOMATRIX embedding medium (CEM) (Thermo Shandon, Pittsburgh, PA) or in Tissue-Tek® optimum cutting temperature (O.C.T.) compound (Sakura Finetek, USA). 5 to 12 µm thick sections were mounted on Super-Frost glass slides (Fisher Scientific) and processed for immunofluorescence. For immunofluorescence labeling, sections were incubated overnight with primary antibody solutions at 4°C in a humidified chamber. After three washes in PBS, sections were incubated with secondary antibodies for 1 h at room temperature. Slides were mounted on coverslips in DAPI-containing mounting medium (Vector Laboratories CA, H-1200). Confocal microscopy analyses were performed using 60x objectives with an IX81 confocal microscope (Olympus, Richmond Hill, Canada), and images were obtained with Fluoview software version 3.1 (Olympus). The cultured cells were fixed for 15 minutes with 4% PFA, washed three times and then permeabilized for 10 min with 0.25 Triton (Sigma, X100-500ML), cells were then blocked in PBS/2% BSA (Sigma, A7906-100G) for an hour and incubated overnight with the primary antibody. Primary antibodies used in this study are rabbit anti-H3K9me3 (1:500, Abcam, ab8898), rabbit anti-H3K9ac (1:500, Cell Signaling, 9671S), rabbit anti-WRN (1:100, Santa Cruz, sc-5629), p-ATR, rabbit anti-53BP1 (1:100, Novus, NB100-304), mouse γH2AX (1:1000, Millipore 05-636), rabbit anti-H2Aub (1:200, Cell Signaling, 8240S), rabbit anti-Ki67 (1:1000, Abcam, ab15580), mouse anti-PCNA (1:250, Invitrogen, MA5-11358), mouse anti-RNAPol2 (1:500, Santa Cruz, sc-47701), and mouse 1H6 antibody recognizing G4s. The 1H6 antibody was obtained from The European Research Institute for the Biology of Ageing. After the primary antibodies, slides were washed three times using PBS and incubated with the secondary antibodies for 1h. The Secondary antibodies are: donkey AlexaFluor488-conjugated anti-mouse (1:1000, Life Technologies), donkey AlexaFluor488-conjugated anti-rabbit (1:1000, Life Technologies), goat AlexaFluor647-conjugated anti-mouse (1:1000, Life Technologies), goat AlexaFluor texas red-conjugated anti-rabbit (1:1000, Life Technologies). Slides were then washed three times with PBS and mounted with coverslips in DAPI-containing mounting medium (Vector Laboratories CA, H-1200).

### Quantifications and statistical analysis

For the colocalization study, random lines were drawn on individual cells using FIJI. From these lines, we plotted the intensity profile of each marker accordingly. The data collected was plotted on horizontal graphs with each marker as a separate line for the visualization of the peaks (example figure 2D). The data was also plotted in a scatter graph, using GraphPad Prism 5, to visualize the correlation between these two markers. On these sets of pairs, a Pearson correlation was calculated to quantify the correlation. For the co-expression study intensity of the signal for different markers was measured, using a mask on DAPI to identify the nucleus, and then plotted in a scatter plot using GraphPad Prism 5, to visualize the correlation between these two markers. On these sets of pairs, a Pearson correlation was calculated to quantify the correlation. For the expression study, we quantified the mean intensity of each marker in the nucleus area using the DAPI signal to identify that area. Values were plotted with a box and whisker graph. The K-means clustering was done searching for 2 or 3 groups with 20 iterations for each run. Statistical differences were analyzed using Student’s t-test for unpaired samples with P ≤ 0.05*, ≤0.01**, ≤0.001***.

### Western Blot

Cell extracts were homogenized in the Complete Mini Protease inhibitor cocktail solution (Roche Diagnostics), followed by sonication. Protein material was quantified using the Bradford reagent. Proteins were resolved in 1x Laemelli reducing buffer by SDS-PAGE electrophoresis and transferred to a Nitrocellulose blotting membrane (Bio-Rad). Subsequently, membranes were blocked for 1h in 5% non-fat milk-1X TBS solution and incubated overnight with primary antibodies. The antibodies used in this study are rabbit anti-BMI1 (Cell Signaling, D42B3, 5856S), mouse anti-LaminA/C (Sant Cruz, E-1, sc-376248), mouse anti-Gapdh (Sant Cruz, G-9, sc-365062). Membranes were then washed 3 times in 1X TBS; 0.05% Tween solution and incubated for 1h with corresponding horseradish peroxidase-conjugated secondary antibodies. Membranes were developed using the Immobilon Western (Millipore).

### Public ChIP-seq analysis and prediction of G-quadruplexes

ChIP-seq datasets were obtained through the GEO platform using accession numbers GSE38273 and GSE33912. BMI1 significant peaks were extracted as in ^34,80^. The Model-based Analysis for ChIP-Seq (MACS) was used to extract significant peaks with a P-value cutoff ≤ 0.05. Peak coordinates were mapped onto hg19 genome reference using SeqMonk v0.34.0 software (Babraham Bioinformatics). Putative G-quadruplexes were predicted using Quadparser algorithm V2 running under python v2.7.11 with indicated parameters for the number of guanines in each stack (G-groups), the number of base pairs between G-groups (loop size) and the number of time the loop and a stack was repeated after the initial stack (Repeats-1)^81^. G-quadruplexes coordinates for each set of parameters were then mapped onto hg19 genome reference using SeqMonk software. Annotation of ChIP-seq peaks with G-quadruplexes was determined by extending them 50 base pairs on each side and counting the number of overlapping predicted G-quadruplexes. SeqMINER was used for H3K9^me3^ ChIP-seq enrichment heatmap and k-means clustering using default parameters.

### Real-time RT-PCR

RNA was isolated using TRIzol reagent (Invitrogen). Reverse transcription (RT) was performed using 1 µg of total RNA and the MML-V reverse transcriptase (Invitrogen). Real-time PCR was carried in triplicates using Platinum SYBRGreen Supermix (Invitrogen) and Real-time PCR apparatus (ABI prism 7002). Primers for *progerin*, *P16INK4A* and *GAPDH* were as in ^59,82^.

## Supplementary Materials

**Figure S1.**
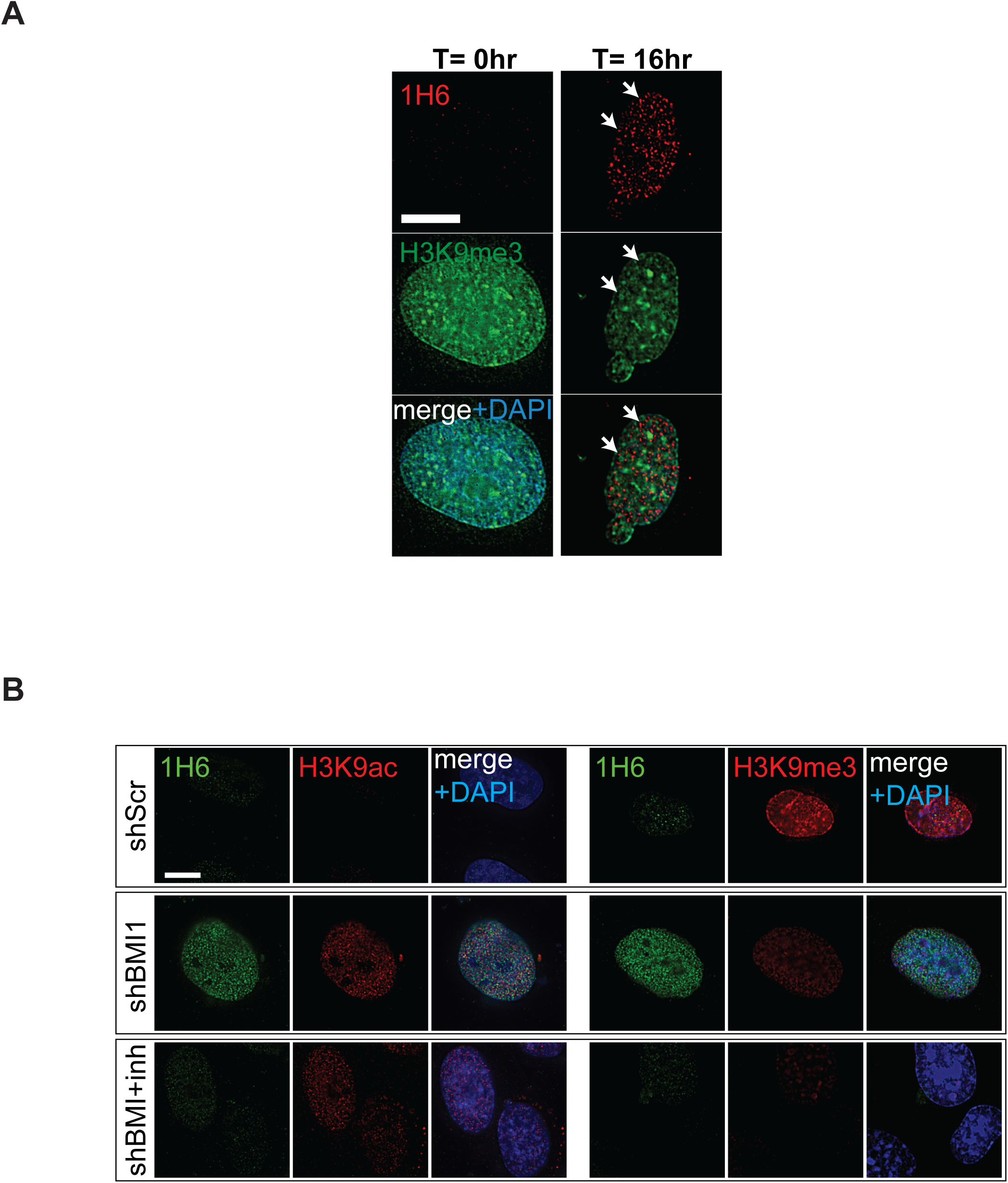
G4 induction inversely correlates with H3K9me3 density. A. Immunofluorescence analysis of shBMI1 HDFs at 0hr and 16hr showing that G4 induction (white arrows) is associated with loss of H3K9me3 density. Scale bar: 5 µm B. Immunofluorescence analysis showing that G4s can be uncoupled from chromatin state, as revealed using transcription (DRB) and replication (Aphi) inhibitors in shBMI1 HDFs. Scale bar: 5 µm

**Figure S2.**
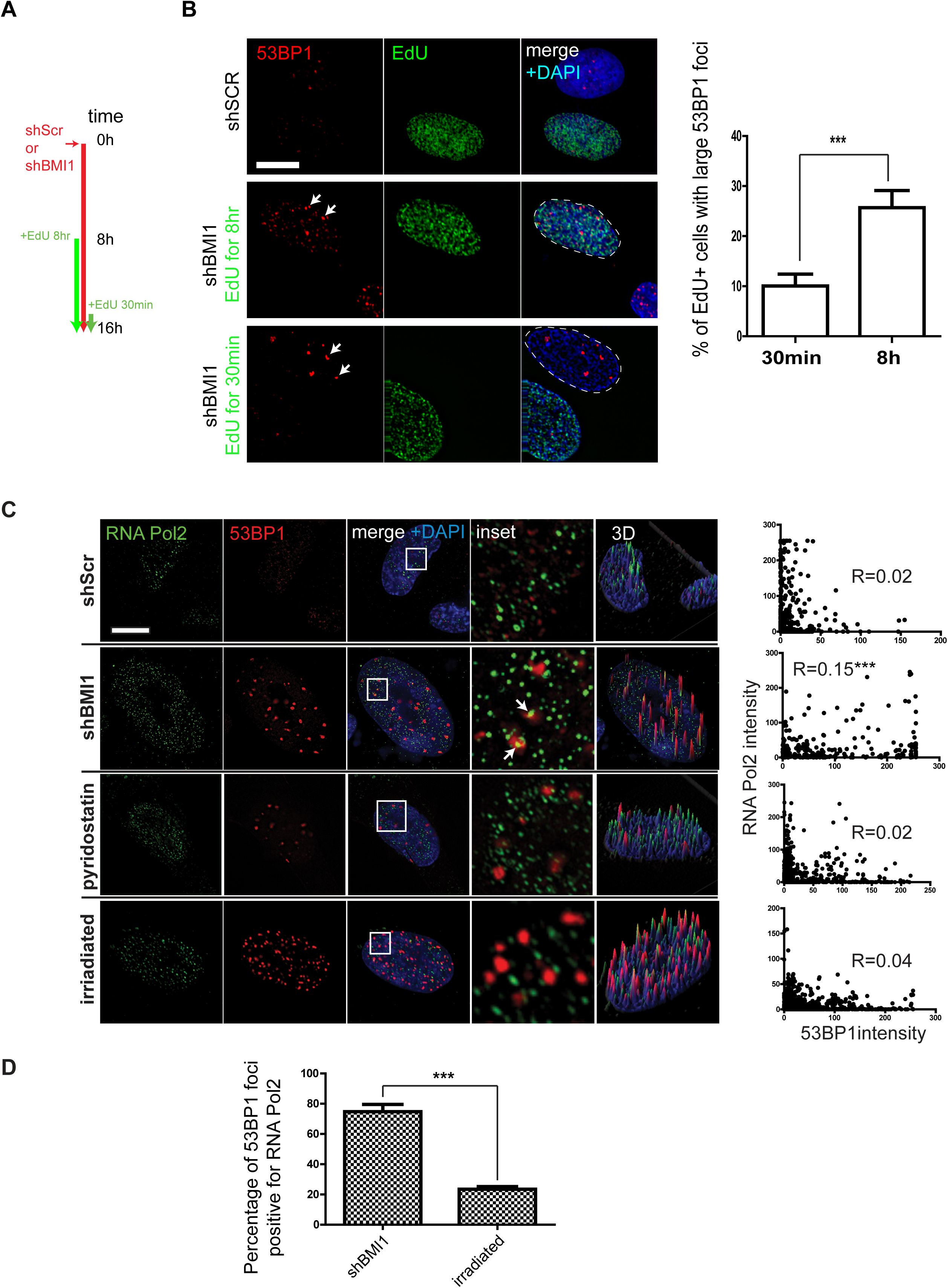
RNA Pol2 localization correlates with DNA damage in shBMI1 HDFs. A. Experimental scheme, where EdU was added at 8hr (for 8 hours) or 15h30hr (for 30min). B. Immunofluorescence analysis of shScr and shBMI1 HDFs exposed to EdU. This showed that the percentage of EdU+ cells with large DDR 53BP1 foci is reduced when cells were labeled at 15h30hr and when compared to 8hr. Arrows indicate large 53BP1 foci in EdU+ and EdU-cells. N = 200 cells/group. Statistical differences were analyzed using an unpaired T-test with two tails. *P* ≤0.001***. All values are means ± SEM. Scale bar: 5 µm C. Immunofluorescence analysis of shScr, shBMI1, or shScr HDFs exposed to pyridostatin for 24 hours or irradiated at 10Gy. Co-localization studies revealed that only shBMI1 HDFs present a correlation between RNA Pol2 and 53BP1 foci (R = 0.15). N = 200 cells/group. Statistical differences were analyzed using an unpaired T-test with two tails. *P* ≤0.001***. Scale bar: 5 µm D. Quantification of experiment in (C) revealing the high percentage of 53BP1 foci that are also positive for RNA Pol2 in shBMI1 HDFs, and when compared to irradiated shScr HDFs. N = 200 cells/group. Statistical differences were analyzed using an unpaired T-test with two tails. *P* ≤0.001***.

